# *Vampyroteuthis pseudoinfernalis* sp. nov.: the second extant widespread deep sea squid species of Vampyromorpha (Cephalopoda: Coleoidea)

**DOI:** 10.1101/2023.02.13.526086

**Authors:** Dajun Qiu, Bilin Liu, Yupei Guo, Wijesooriya A.S.W. Lakmini, Yehui Tan, Gang Li, Zhixin Ke, Kaizhi Li, Liangmin Huang

## Abstract

The vampire squid, *Vampyroteuthis infernalis* Chun, 1903, is currently the only extant species in the family Vampyroteuthidae Chun, 1903. However, specimens from the Gulf of Guinea, Africa, and California suggested the possibility of additional taxa. Here, we report the second species of *Vampyroteuthis*, collected from the South China Sea, China, which differs from *V. infernalis* by the tail shape, the lower beak, and the phylogenetic analysis of mitochondrial COI and nuclear large subunit ribosomal DNA (LSU) sequences: *V. infernalis* present by the lacking of the tail, photophores located approximately one-third of the points between the fins and end, and the lower beak with a broad, short wing; *V. pseudoinfernalis* Qiu, Liu & Huang, **sp. nov.** present by an acuate tail, a pair of photophores located at the midpoints between the fins and tail, and a lower beak with a broad, elongate wing.

## 1 Introduction

Planktonic cephalopods are widely distributed in the world’s oceans from the surface to abyssal depths, where they occupy epipelagic, mesopelagic and bathypelagic zones. One of these cephalopods, *Vampyroteuthis infernalis* Chun, 1903, is currently the only extant species in Vampyroteuthidae. The species was initially described by Chun (1903) under the order Octopoda (Chun, 1903; Pickford, 1946). Then, at least seven genera and ten species were reported by different authors (Robson, 1932). Pickford (1939) reclassified the eleven extant species as a single species, *V. infernalis*, and transferred it from the order Octopoda to the new order Vampyromorpha (Pickford, 1939, 1946).

The recognized distribution of *V. infernalis* includes temperate and tropical Pacific, Indian, and Atlantic oceans (Yokobori *et al*., 2007; Jereb *et al*., 2010; Hoving & Robison, 2012), where it typically occurs between 600 m and 900 m at oxygen concentrations of *ca.* 0.4 mL/L in the northeastern Pacific (Hoving & Robison, 2012). However, morphological differences had been reported among specimens from the Gulf of Guinea, Africa, and California (Young, 1972), and genetic differences or population structure analysis suggest that two or multiple extant species may exist in this family (Braid & Bolstad, 2019; Timm *et al*., 2020).

The family Vampyroteuthidae and the genus *Vampyroteuthis* was not previously reported from the South China Sea, where presents 153 cephalopod species (Norman *et al*., 2016). Here we describe a new species, namely *Vampyroteuthis pseudoinfernalis* Qiu, Liu & Huang, **sp. nov.**, with its small (SSU) and large (LSU) rDNA and mitochondrial cytochrome c oxidase subunit I (COI) sequences.

## 2 Materials and methods

### 2.1 Sample collection, research ethics, and observations

The specimen was collected during an open cruise near Hainan Island, northwestern South China Sea (Stn 30: 17°59*’*28*”*N, 112°29*’*58*”*E), on 13 September 2016 (Fig. 1) from a depth between 800 m and 1000 m depth using a Hydro-Bios Multinet system (Type Max, mouth area 0.5 m^2^, 300 µm mesh; HydroBios Inc., Kiel, Germany) hauled vertically at approximately 0.5 m/s. The temperature at this station ranged 5.75–8.86°C, and the salinity ranged 34.46–34.36 PSU. The net was equipped with internal and external flow meters and a depth transducer, all of which were monitored and logged in real time; sampling of the water column otherwise occurred in 1000–800 m, 800–600 m, 600–400 m, 400–300 m, 300–200 m, 200–100 m, 100–50 m, 50–25 m and 25–0 m.

**Figure 1.**
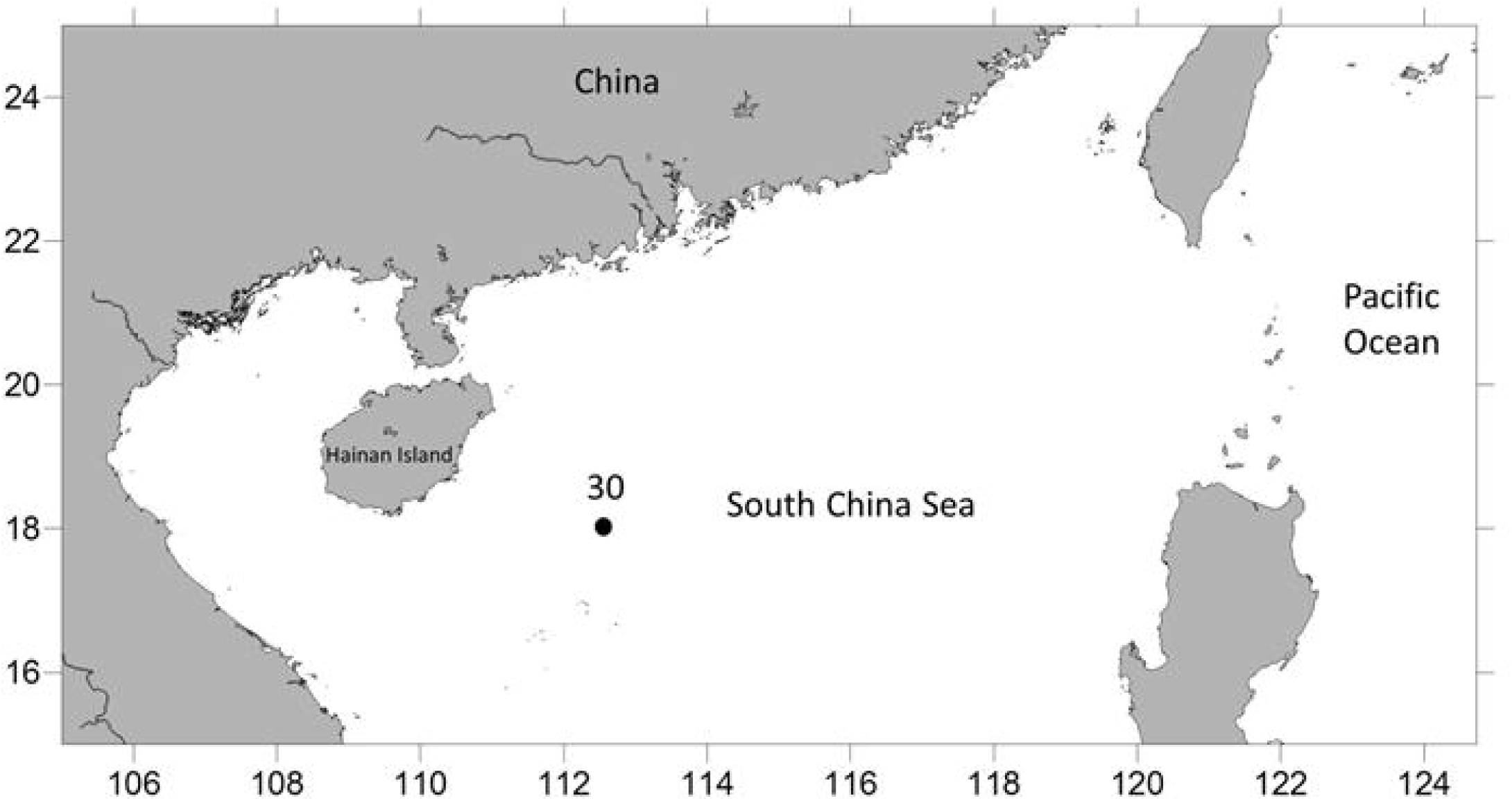
Map of study site. The *Vampyroteuthis* was collected from Station 30.

Immediately after sorting, the specimen was photographed, and counts and measures were taken following Roper and Voss (1983). The specimen was then frozen at -20°C and kept in darkness for subsequent molecular and morphological analyses, deposited in the Marine Biodiversity Collection of South China Sea, South China Sea Institution of Oceanology, Chinese Academy of Sciences.

### 2.2 DNA extraction, PCR, and gene sequencing

A tissue sample was resuspended in 0.5 mL of DNA lysis buffer (0.1 MEDTA pH 8.0, 1% SDS, 200 µg/mL proteinase K) in a 1.5 mL tube and then incubated for 48 hours at 55°C. DNA extraction and purification were performed according to the protocols of Qiu *et al*. (2011). After extraction, the sample DNA was eluted in 50 µL of Tris-HCl solution before 1 µL of DNA was subjected to PCR with pairs of primers (SSU, LSU, and COI) to amplify the SSU rDNA, LSU rDNA, and mitochondrial COI genes, respectively. Primer sequences are detailed in Table 1.

**Table 1.**
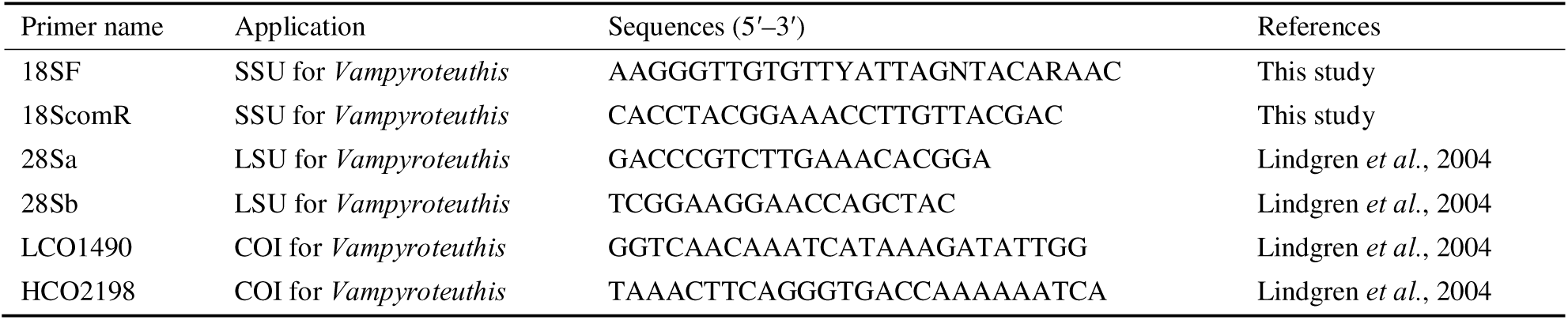
Primers used in this study.

PCR amplification was carried out in 25 µL reaction volumes under thermocycling conditions, including a denaturing step of 94°C for 4 min; 35 cycles of 94°C for 30 sec, 56°C for 30 sec, and 72°C for 45 sec; and a final extension step of 72°C for 10 min. PCR products were resolved by agarose gel electrophoresis with a DL2000 DNA Ladder (TaKaRa Bio, Dalian, China) before bands of the expected sizes were excised to remove primer dimers. The DNA was purified and directly sequenced as described by Qiu *et al*. (2011).

### 2.3 Phylogenetic analyses

The sequences were trimmed with primers, and the two strands were merged. The assembled sequences were analyzed against the GenBank database using the Basic Local Search Tool (BLAST). Sequences showing significant similarity in BLAST to our sequences were downloaded. Phylogenetic analysis based on partial SSU (1899 bp), partial LSU (D1-D2, 408 bp), and mitochondrial COI (702 bp) regions was done to investigate the phylogenetic position of *Vampyroteuthis* specimens. The combined sequences were aligned with ClustalW. The alignment was run through ModelTest to select the most appropriate evolutionary model. The selected general time-reversible model with a gamma distribution was subjected to maximum likelihood analysis using MEGA 5.0. Categories of substitution rates were set at 4, and other parameters were estimated based on the datasets. The sequences obtained in this study were deposited in GenBank under accession numbers MN044751, OP526744, OP531921, and OP549780.

## 3 Results

The result strongly supports the existence of two distinct *Vampyroteuthis* species which differ in tail shape, photophore position, lower beak morphology, and evolutionary distance between the respective LSU rDNA and mitochondrial COI sequences. We propose that one of these taxa, the South China Sea specimen, represents a new species, namely *V. pseudoinfernalis* Qiu, Liu & Huang, **sp. nov.**

### 3.1 Taxonomy

Vampyroteuthis pseudoinfernalis Qiu, Liu & Huang, sp. nov.

#### Diagnosis

The new species is different from its sibling species *V. infernalis* by with an acuate tail. It is characterised by the following combination of morphological characters: a pair of photophores located at the midpoints between the fins and tail, and a lower beak with a broad, elongate wing.

#### Description

Mature female specimen (Fig. 2) small, extensively gelatinous, with black and reddish-brown chromatophores interspersed over body surfaces. Total length 9.5 cm, mantle length 5.2 cm and width 5.0 cm. Mantle conical and rounded, with an acuate tail (Fig. 2). Head fused with mantle; brachial and nuchal constrictions poorly demarcated. Eyes brown (0.7 cm diameter), recessed into head tissues, and covered by a crystalline membrane. Arms with comparable length to mantle, subequal, with tapering ends. Web depth *ca.* 2.5 cm, *ca.* 50% arm length. Suckers small, in single series, on distal half of arms only, unstalked, without cuticular lining. Two rows of cirri extend along each arm for length. Threadlike arm ends naked at medial of it and web margin. A pair of long and paddle-shaped fins (width *ca.* 5 cm, length *ca.* 0.9 cm), lacking a cartilaginous base, protruded from posterior mantle. A pair of large photophores (*ca.* 0.3 cm diameter) lies between fins and tail. Distances from fins to photophores and from photophores to tail were 1.0 cm, 1.0 cm, respectively. All distance data are presented in Table 2.

**Figure 2.**
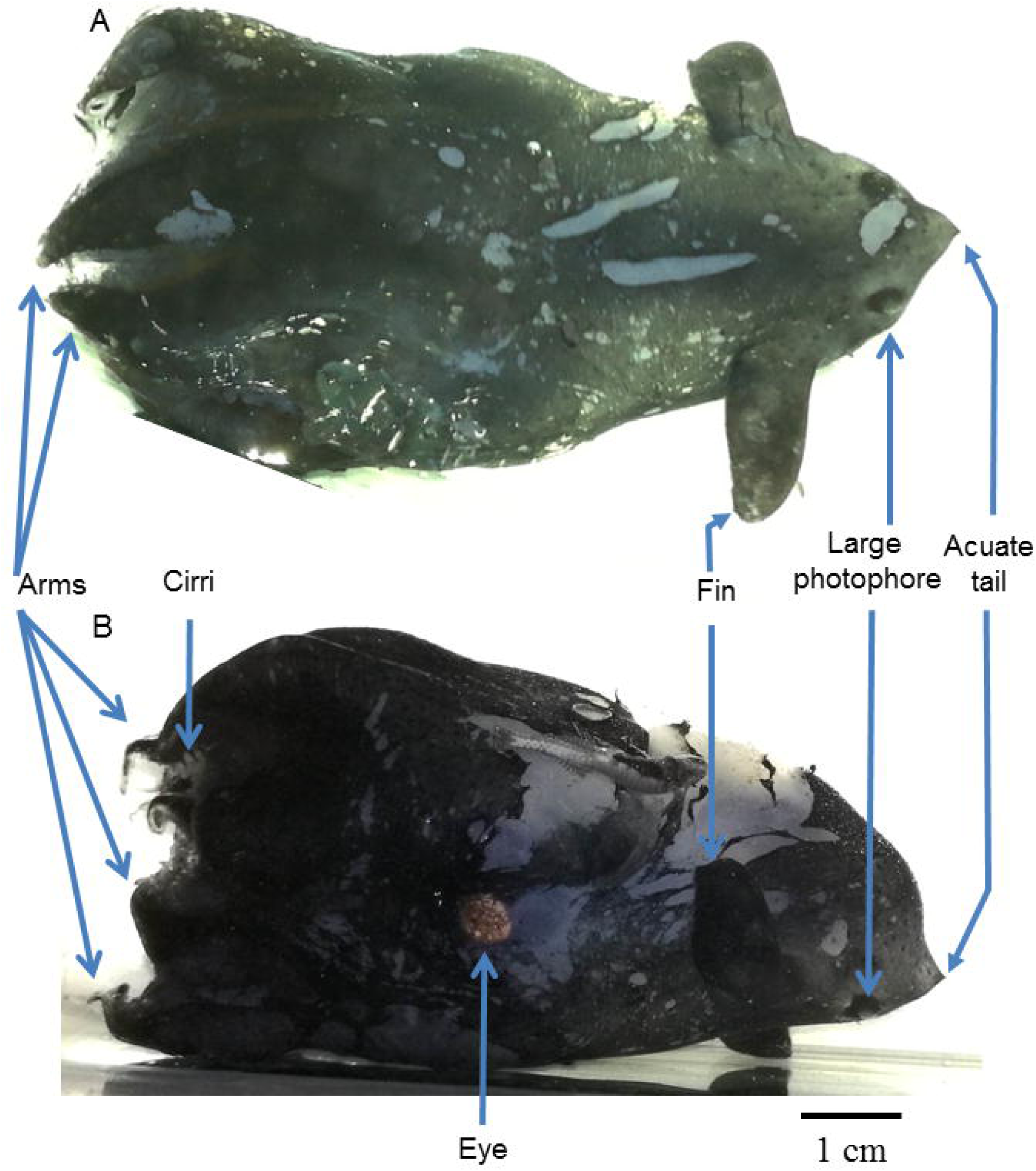
Fresh-collected *Vampyroteuthis pseudoinfernalis* Qiu, Liu & Huang, sp. nov.: A) dorsal, B) lateral views. Scale bar = 1 cm.

**Table 2.** Distances from the fins to the photophores and from the photophores to the tail or end of Vampyroteuthis.

| Specimen number | Species | From fins to photophores (cm) | From photophores to tail or end (cm) | Reference |
| --- | --- | --- | --- | --- |
| 1 | *Vampyroteuthis pseudoinfernalis* | 1.0 | 1.0 | This study |
| 2 | *V. infernalis* | 0.6 | 1.15 | Pickford, 1949: plate II, fig. 4. |
| 3 | *V. infernalis* | 0.2 | 0.4 | Pickford, 1949: plate III, fig. 8. |
| 4 | *V. infernalis* | 0.3 | 0.7 | Pickford, 1949: plate III, fig. 9. |
| 5 | *V. infernalis* | 0.15 | 0.3 | Pickford, 1949: plate III, fig. 7. |

#### Gladius

Gladius broad, chitinous, and transparent, covering dorsal mantle beneath skin and a thick layer of loose connective tissue (Fig. 3). Gladius thin and flat in anterior, gradually thickening and arching strongly in posterior. Gladius consists of middle (ostracum), inner (hypostracum) and outer (periostracum) shell layers. Proostracum comprises five longi-tudinal elements: a middle plate (rachis), two lateral plates, and two wings (Fig. 3A).

**Figure 3.**
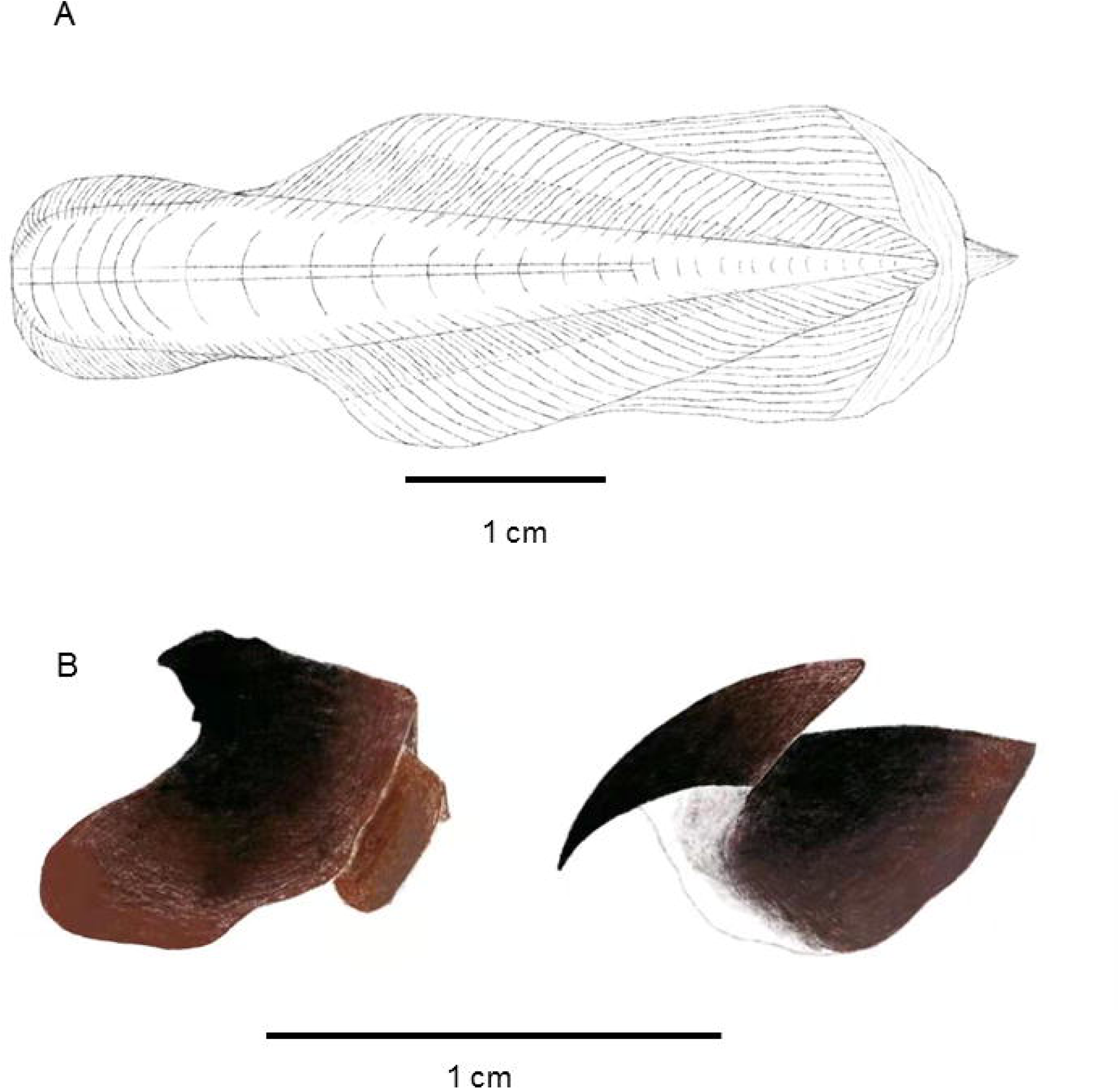
Gladius (A), upper and lower beak (B) of *Vampyroteuthis pseudoinfernalis* Qiu, Liu & Huang, sp. nov. Scale bar = 1 cm.

#### Beaks

Upper beak rostrum long, curved, tip pointed. Jaw angle obtuse, lateral wall extends forward to wing, where becomes large and with a distinct false angle. Hood long, *ca.* 0.8 times of crest length. Posterior margins of hood/wing convex. Wing extends to base of anterior margin of lateral wall. Crest straight and unthickened (Fig. 3B). Lower beak rostral tip pointed, with a small hook. Wings very broad, elongate, and spread parallelly, with a very high wing fold and highest opposite jaw angle, forming a smooth cutting edge. Crest short, wide, unthickened. Jaw angle obtuse. Shoulder tooth present (Fig. 3B). Beak measurements presented in Table 3.

**Table 3.**
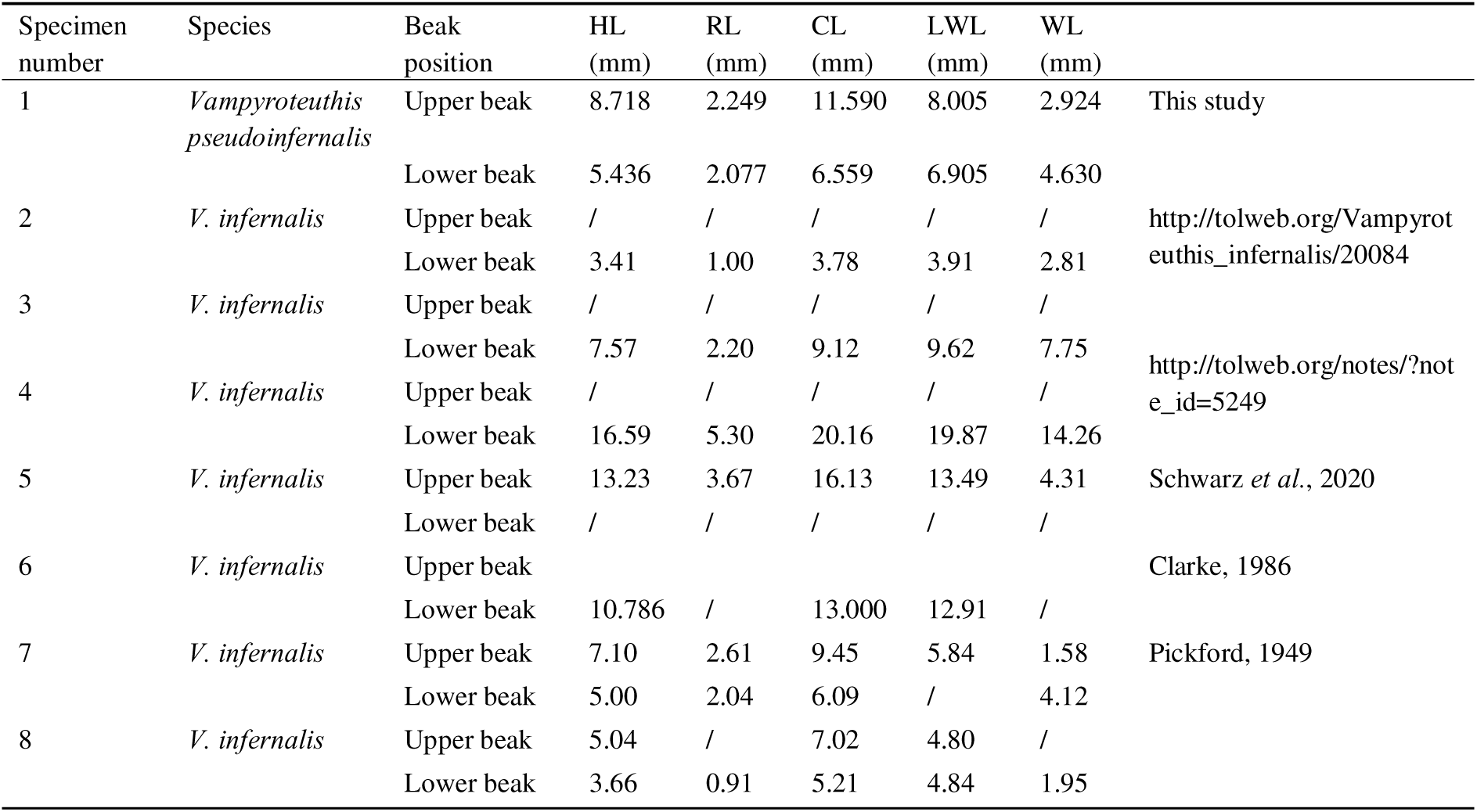
Lengths of the upper and lower beak characters of *Vampyroteuthis* in this study and literature data. HL—hood length; RL—rostral length; CL—crest length; LWL—lower wing length; WL—wing length.

#### Holotype

Hainan Island, northwestern South China Sea (Stn 30: 17°59*’*28*”*N, 112°29*’*58*”*E), 13 September 2016, coll. Dajun Qiu (SCSMBC030991). The specimen is barcoded in GenBank as follows: SSU 1-1899 (MN044751), D1-D2 LSU 1-408 rDNA (OP526744), and COI 1-702 (OP531921) sequences.

#### Etymology

The specific name is derived from the Latin word “*pseudo*” and the previous species *V. infernalis*.

#### Distribution

The species is known only in the South China Sea now. However, the molecular data shows it has a wider distribution, in the Gulf of Mexico, Atlantic Ocean, Cape Verde, northern Atlantic Ocean, off Tohoku (North Pacific Ocean), and western Pacific Ocean (D1–D2 LSU rDNA: MG263928, MG263931, MG263941, MG263942, MG263948, MG263949, MG263954, MG263969, MG263987, MG263992; COI: AB266515, AB385880, GU145060, MF432197, OR378923, OR379049, MG591300, MG591304, MG591313, MG591328, MG591329, MG591335, MG591338, MG591339, MG591358, MG591379, MG591380, MG591385, MG591386, MG591404, MG591438, MG591440, MG490128, NC009689; data from GenBank).

### 3.2 Phylogenetic position of *Vampyroteuthis* based on nuclear rRNA and mitochondrial COI genes

Nuclear-encoded partial SSU (MN044751) and LSU (OP526744) ribosomal RNA genes and mitochondrial COI (OP531921) were obtained. The phylogenetic tree of SSU, LSU, and COI contain 40, 29, and 80 sequences, respectively, from GenBank, besides our sequences. The LSU tree topologies inferred from the dataset using neighbor joining (NJ) and maximum likelihood (ML) revealed well-supported long distances and clear separation of *V. pseudoinfernalis* Qiu, Liu & Huang, **sp. nov.** from *V. infernalis* and other genera (Fig. 4). Because our LSU rRNA gene sequence is almost identical to 10 sequences (MG263928, MG263931, MG263941, MG263942, MG263948, MG263949, MG263954, MG263969, MG263987, MG263992) from the Gulf of Mexico and Atlantic Ocean, they may be conspecific. The LSU phylogeny clusters *V. pseudoinfernalis* Qiu, Liu & Huang, **sp. nov.** in a distinct lineage separated from 14 other sequences of *V. infernalis* from the Gulf of Mexico (MG263894, MG263895, MG263896, MG263897, MG263926, MG263930, MG263940, MG263951, MG263953, MG263959, MG263985, MG263986) and two unknown localities (AH012197 and AY557548) (Fig. 4).

**Figure 4.**
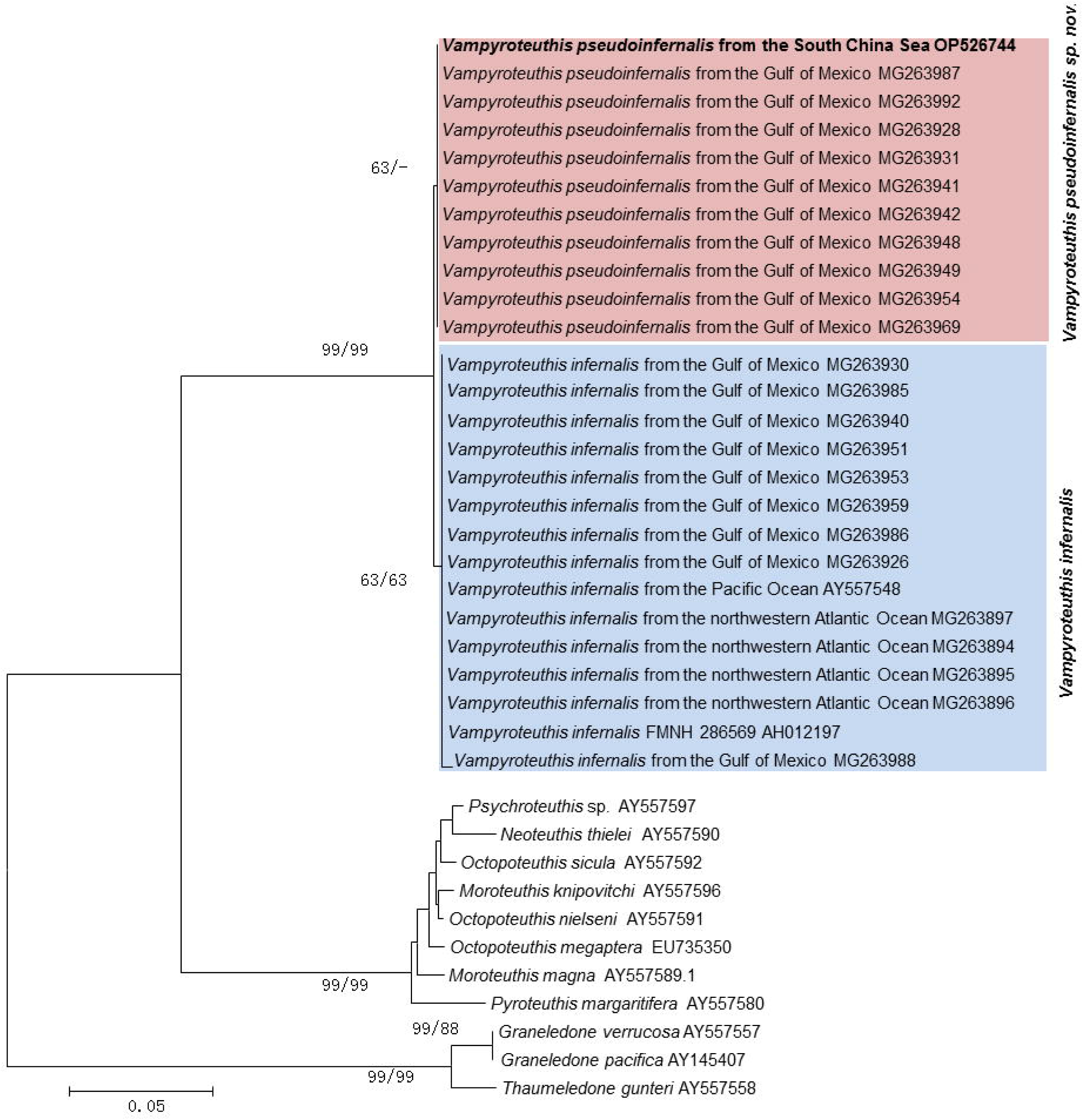
Cephalopod phylogeny inferred from LSU rDNA. Nodal support based on bootstrap values of ML/NJ with 1000 resamplings each; only values > 60 are shown (if a phylogenetic method yielded support < 60, it is denoted by ‘‘-’’). *Thaumeledone gunteri* was used as an outgroup to root the tree.

In the mitochondrial COI tree (Fig. 5), the South China Sea sequence is identical to the 21 sequences reported for *V. infernalis* from the Gulf of Mexico, the northeast and northwest Atlantic Ocean, and the North Pacific Ocean (off Tohoku) and forms a strongly supported distinct lineage that is well separated from 20 sequences referred to *V. infernalis* from the Gulf of Mexico, near New Zealand (South Pacific Ocean), the Indian Ocean, and the northwest Atlantic Ocean. Thus, the mitochondrial COI tree divided them into two groups. For COI, a total of 8 diagnostic positions were identified between the two groups (Table 4, Fig. S1). The sequences of AY557459 (SSU rDNA, Fig. 6), AY557548 (LSU rDNA, Fig. 4), and AF000071 (COI, Fig. 5) were sourced from the same specimen referred to as *V. infernalis* by Carlini *et al*. (2001) and Lindgren *et al*. (2004). The sequence of MK186003 (COI, Fig. 5) clustered with the sequence AF000071 in a group that was sourced from New Zealand specimens and referred to as *V. infernalis*. In the SSU tree (Fig. 6), the 1899-bp SSU rDNA sequence from the South China Sea of *V. pseudoinfernalis* Qiu, Liu & Huang, **sp. nov.** (MN044751) differed by only 5 bp (0.3%) from that of *V. infernalis* (AY557459, AY145387). Therefore, the SSU rDNA sequence of *V. pseudoinfernalis* Qiu, Liu & Huang, **sp. nov.** could be difficultly separated from the *V. infernalis* sequence (AY557459) based on the sequence. As a result, the molecular phylogenies revealed that *V. pseudoinfernalis* Qiu, Liu & Huang, **sp. nov.** formed strongly supported lineages (Figs 4–6).

**Figure 5.**
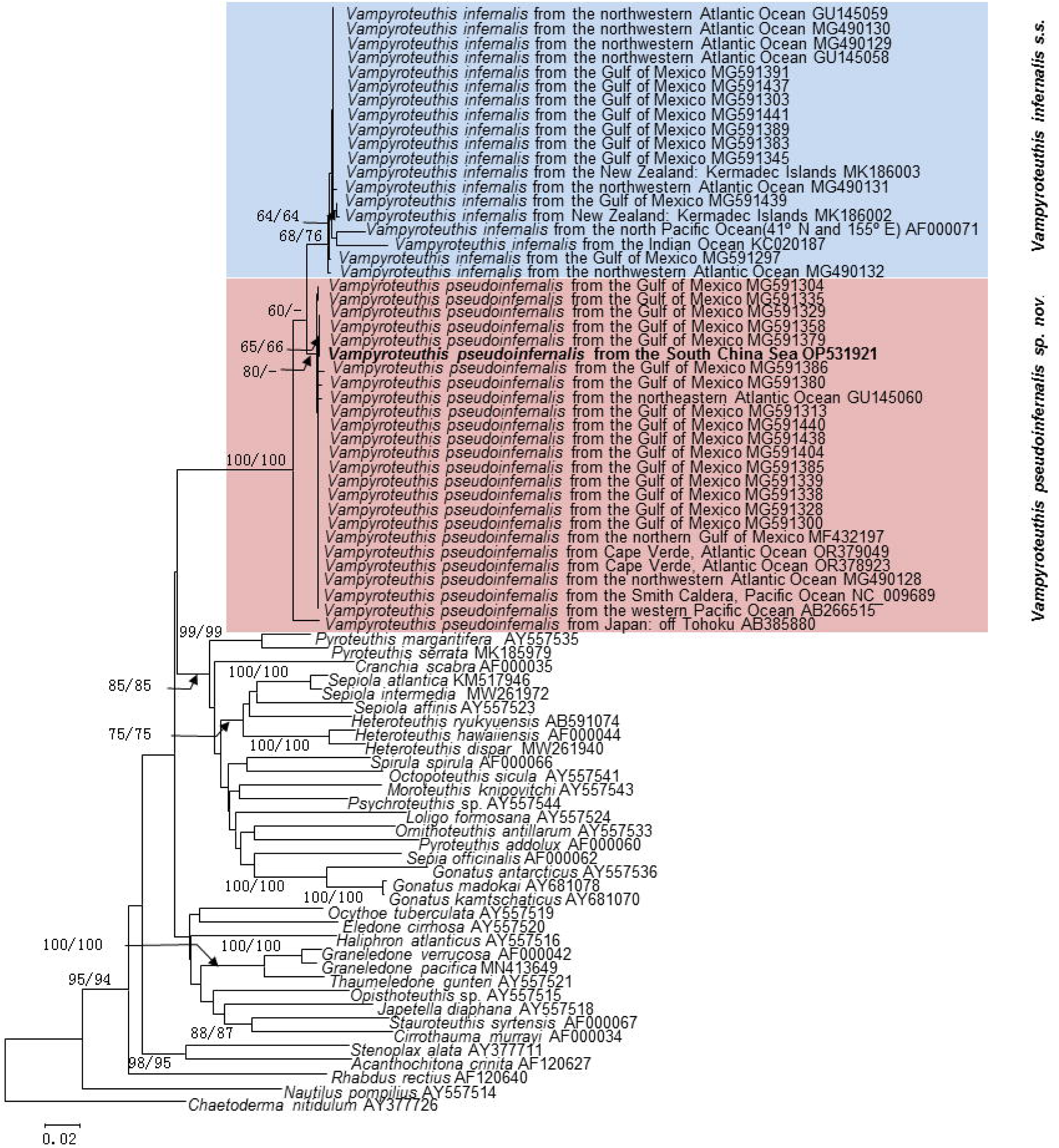
Cephalopod phylogeny inferred from mitochondrial COI. Nodal support based on bootstrap values of ML/NJ with 1000 resamplings each; only values > 60 are shown (if a phylogenetic method yielded support < 60, it is denoted by ‘‘-’’). *Chaetoderma nitidulum* was used as an outgroup to root the tree.

**Figure 6.**
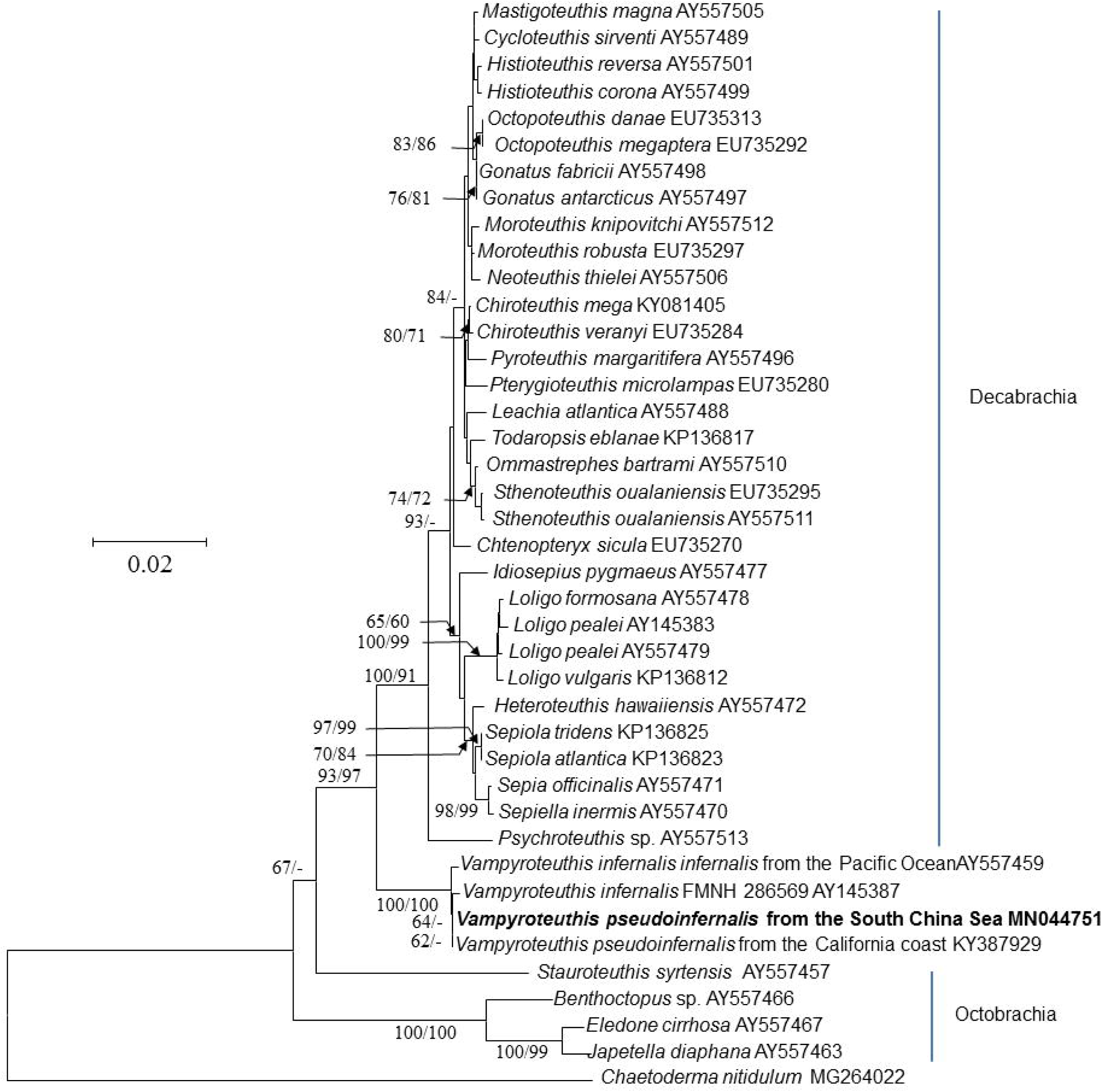
Cephalopod phylogeny inferred from SSU rDNA. Nodal support based on bootstrap values of ML/NJ with 1000 resamplings each; only values > 60 are shown (if a phylogenetic method yielded support < 60, it is denoted by ‘‘-’’). *Chaetoderma nitidulum* was used as an outgroup to root the tree.

**Table 4.** Molecular diagnostic characters obtained from cytochrome c oxidase subunit I (COI)

| Position | *Vampyroteuthis infernalis* | *V. pseudoinfernalis* sp. nov. | *Gonatus madokai* | *Graneledone verrucosa* |
| --- | --- | --- | --- | --- |
| 72 | G | A | A | A |
| 180 | G | A | G | A |
| 372 | G | A | C | T |
| 474 | A | T | A | A |
| 517 | C | T | T | T |
| 528 | A | T | A | A |
| 618 | A | T | G | T |
| 639 | C | T | T | T |

The alignment is provided in Supporting FASTA File S1 to name each nucleotide position unambiguously.

## 4 Discussion

### 4.1. Vampyroteuthis infernalis sensu lato

Molecular phylogeny indicates that *Vampyroteuthis infernalis sensu lato* is strongly bootstrap supported to be divided into two separated species, *V. pseudoinfernalis* Qiu, Liu & Huang, **sp. nov.** and *V. infernalis sensu stricto* (Figs 4–6, Table 4). In morphology, these two species can also be distinguished by their tail shape, lower beak, and photophore position.

The species *V. infernalis* was originally described by Chun (1903), however, he did not record the position of photophore (Fig. 7A). Similar situations are also present in its four synonyms, *Cirroteuthis macrope*, *Melanoteuthis beebei*, *Watasella nigra* and *Danateuthis schmidti* (Figs 7B, G–I; Chun, 1903; Berry, 1911; Sasaki, 1920; Joubin, 1929; Robson, 1929; Pickford, 1949). Another five synonyms have the original description of the photophore position, such as *Retroteuthis pacifica* (Figs 7C), *Hansenoteuthis lucens* (Figs 7E–F), *Melanoteuthis lucens* (Fig. 7H), *Melanoteuthis schmidti* (Figs 7J–K), and *Melanoteuthis anderseni* (Figs 7I–M) (Joubin, 1912, 1929, 1931; Pickford, 1949). *Retroteuthis pacifica* showed the distortion of tail shape which did not present the true photophores positions (Fig. 7C-D; Joubin, 1929). Based on these descriptions/monographs, the ratio of the distance ‘from fins to photophores’ to the distance ‘from photophores to tail’ is far less than 1 (*Retroteuthis pacifica*) or approximately 0.5 (juvenile individuals of *Hansenoteuthis lucens* and *Melanoteuthis schmidti*, and mature individuals of *Melanoteuthis anderseni*). In this study, the corresponding ratio is 1 (Tables 2, 5, Fig. 7). Additionally, all monographs of *V. infernalis* and its synonyms not recorded the tail.

**Figure 7.**
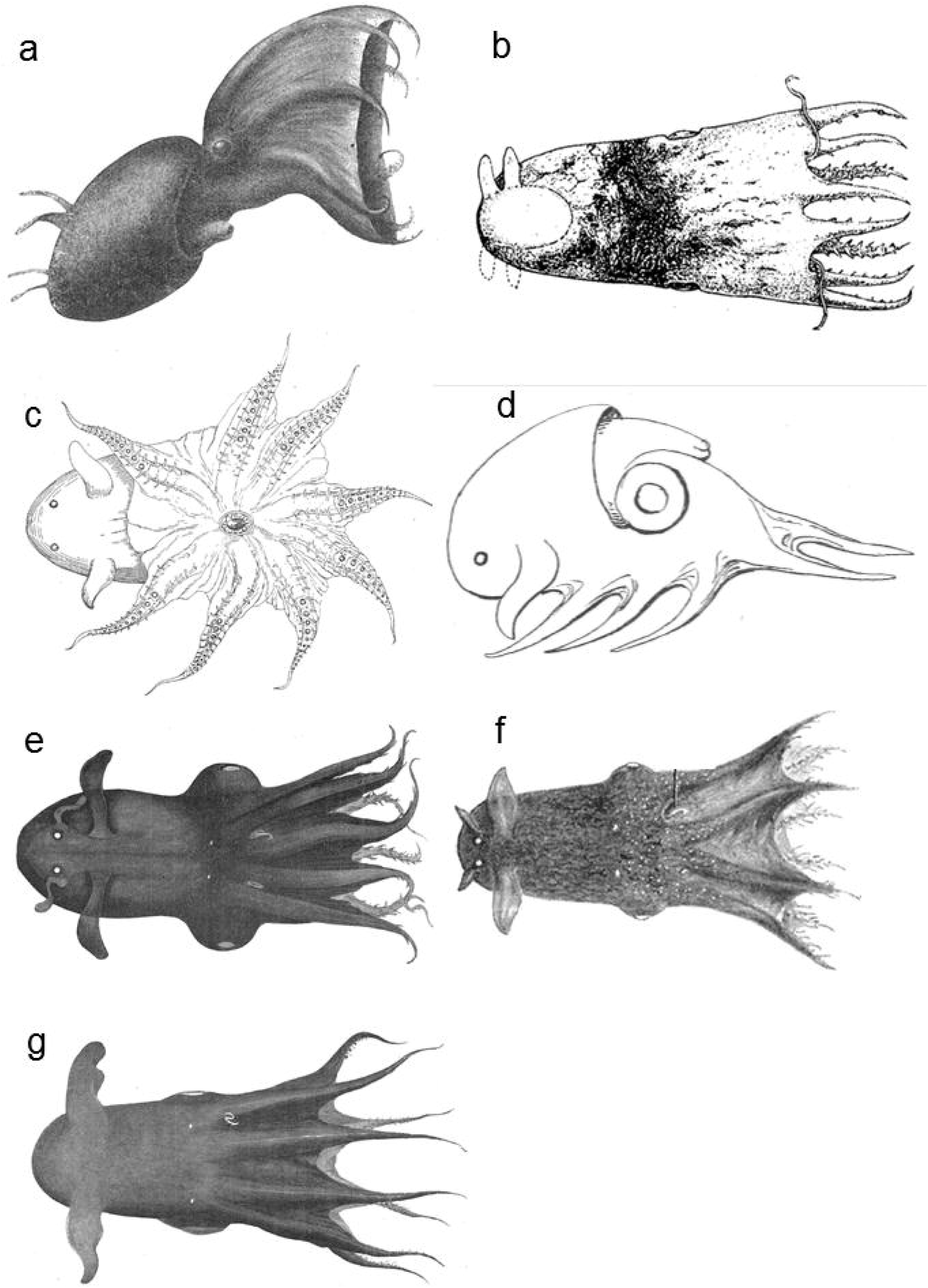
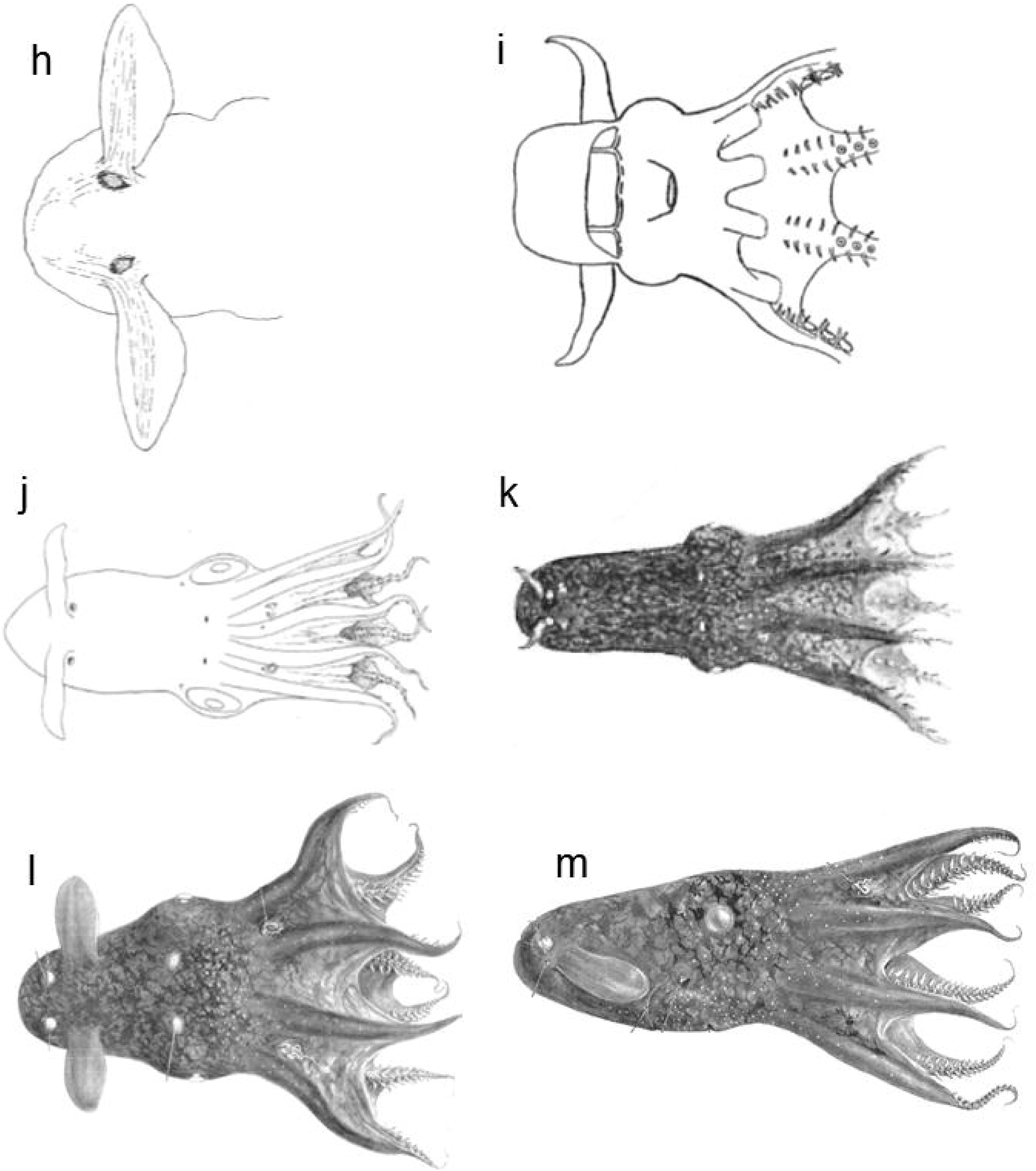
Drawings of *Vampyroteuthis infernalis*, *Melanoteuthis* spp., *Watasella nigra*, *Danateuthis schmidti*, *Retroteuthis pacifica*, and *Hansenoteuthis lucens* from the literature. (a) Lateral view of *Vampyroteuthis infernalis* from Chun (1903). (b) Dorsal view of *Watasella nigra* from Sasaki (1920). (c-d) Dorsal and lateral view of *Retroteuthis pacifica* from Jubin (1929, Fig.13, 15). (e) Dorsal view of *Hansenoteuthis lucens* from Jubin (1929, Fig.17). (f) Dorsal view of *Hansenoteuthis lucens* from Pickford (1949, Fig. 9). (g) Dorsal view of *Danateuthis schmidti* from Jubin (1929, Fig.7). (h) Dorsal view of *Melanoteuthis lucens* from Jubin (1912, Fig. 2). (i) Outline drawing of *Melanoteuthis beebei* (ventral view, partly restored) from Roboson (1929). (j) Dorsal view of *Melanoteuthis schmidti* from Jubin (1929, plate III, Fig.3). (k) Dorsal view of *Melanoteuthis schmidti* from Pickford (1949, plate III, Fig.2, 4). (l-m) Dorsal and lateral view of *Melanoteuthis anderseni* from Pickford (1949, plate II, Fig. 4; plate I, Fig. 1).

**Table 5.**
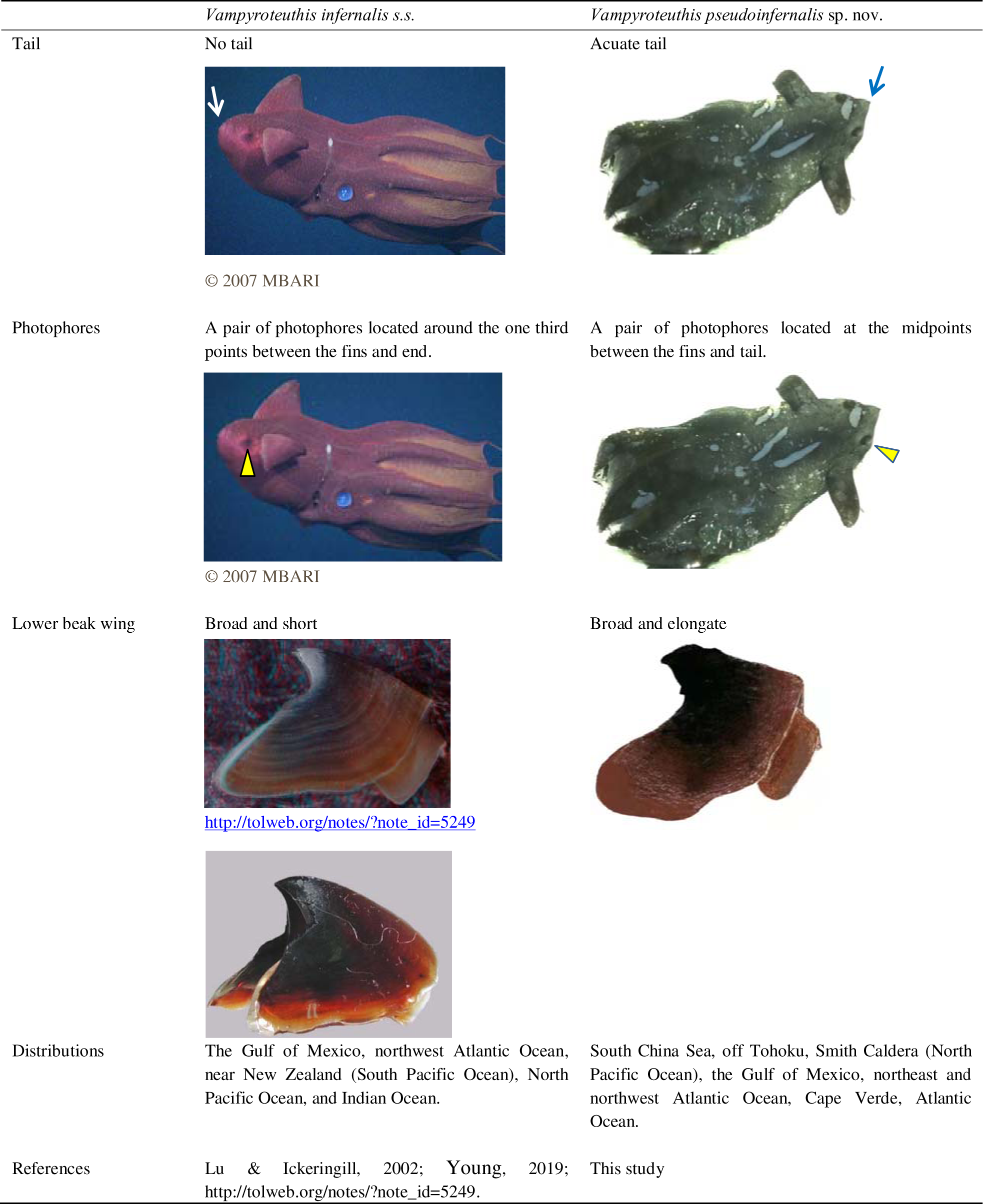
Morphological characteristics and distributions of *Vampyroteuthis* species.

|  | *Vampyroteuthis infernalis* s.s. | *Vampyroteuthis pseudoinfernalis* sp. nov. |
| --- | --- | --- |
| Tail | No tail<br>© 2007 MBARI | Acuate tail |
| Photophores | A pair of photophores located around the one third points between the fins and end.<br>© 2007 MBARI | A pair of photophores located at the midpoints between the fins and tail. |
| Lower beak wing | Broad and short<br>http://tolweb.org/notes/?note_id=5249 | Broad and elongate |
| Distributions | The Gulf of Mexico, northwest Atlantic Ocean, near New Zealand (South Pacific Ocean), North Pacific Ocean, and Indian Ocean. | South China Sea, off Tohoku, Smith Caldera (North Pacific Ocean), the Gulf of Mexico, northeast and northwest Atlantic Ocean, Cape Verde, Atlantic Ocean. |
| References | Lu & Ickeringill, 2002; Young, 2019; http://tolweb.org/notes/?note_id=5249. | This study |

The new species is different from the *V. infernalis* populations of the southeastern and northeastern Pacific and central and eastern Atlantic (Clarke, 1986; Lu & Ickeringill, 2002; Schwarz *et al*., 2020) by having a broad and elongated wing of lower beak. Its ratio of lower beak crest length (CL) to hood length (HL) is similar to that of previous studies, except specimen 8 (Table 3). The ratio of lower beak wing length (WL) to rostral length (RL) is significantly differed from specimen 2, 3 and 4, but was similar to specimen 7 and 8 (Table 3).

In conclusion, *V. pseudoinfernalis* Qiu, Liu & Huang, **sp. nov.** is characterized by having an acuate tail, photophores located at the midpoints between the fins and tail, and a broad and elongated wing of lower beak (Tables 2, 5), while *V. infernalis sensu stricto* has no recorded tail, photophores located about one-third of the fins and end, and the lower beak with a broad and short wing.

According to the molecular phylogenies, *V. pseudoinfernalis* Qiu, Liu & Huang, **sp. nov.** branched at a basal position to *V. infernali*s *sensu stricto*; these two species indisputably share common ancestry (Figs 4–5). Timm *et al*. (2020) reported that evidence of two populations of *Vampyroteuthis* was found in the Gulf of Mexico and northwestern Atlantic Ocean based on population structure analysis.

### 4.2 Phylogenetic position of Vampyromorpha

The phylogenetic analyses support the point that Vampyromorpha occupies an intermediate phylogenetic position between Decabrachia and Octobrachia (Fig. 6). Among the surveyed taxa, Vampyromorpha appears to be more closely related to *Psychroteuthis* sp. (AY557513, differing in 345 bp (15.92%)) than to *Stauroteuthis syrtensis* (differing in 851 bp (32.78%)). The result is similar to the views of Lindgren *et al*. (2004), Pickford (1939, 1940), Young (1977), Healy (1990), Young & Vecchione (1996) and Young *et al*. (1998), that Vampyromorpha was more closely related to Decabrachia (or Teuthida) than Octobrachia (particularly cirrate octopuses) based on either molecular or morphological data. On the contrast, Yokobori *et al*. (2007) suggested than Vampyromorpha was more closely related to Octobrachia based on mitochondrial gene sequence information.

However, our results and historical data have yet to unequivocally demonstrate the monophyly of Decabrachia and Vampyromorpha or Octobrachia and Vampyromorpha. Further analyses are needed to determine the phylogenetic relationships between these taxa.

## Authorship contribution

Dajun Qiu: Conceptualization, Investigation, Methodology, Data curation, Formal analysis, Writing – original draft, Writing – review & editing, Project administration, Funding acquisition. Bilin Liu: Methodology, Formal analysis, Writing – original draft, Writing – review & editing. Liangmin Huang: Supervision, Project administration, Funding acquisition. Yupei Guo: Formal analysis. W.A.S.W Lakmini: Formal analysis. Yehui Tan: Analysis tools. Gang Li: Investigation. Zhixin Ke: Investigation. Kaizhi Li: Analysis tools.

## Acknowledgements

The authors thank Guozheng Wang, Huangchen Zhang, Yanan Tang, Yu Zhong, and Chenhui Xiang from the South China Sea Institute of Oceanology, Chinese Academy of Sciences, and colleagues of the northern South China Sea Marine Research Open Cruises (NSFC) for helping us use the Hydro-Bios Multinet system or the collection of deep-sea samples. We thank Dr. Fuqiang Chen and two reviewers for help with English language and comments. We thank Richard Young from the University of Hawaii for providing and permissioning the *Vampyroteuthis infernalis* beak photo and insightful advice on the data analysis. We also thank the Monterey Bay Aquarium Research Institute (MBARI) for providing and permissioning a *Vampyroteuthis infernalis* digital image.

## References

Berry, S.S. 1911. Preliminary Notices of some new Pacific Cephalopods. Proceedings of the United States National Museum, 40(1838): 589–592.

Braid, H.E., Bolstad, K.S.R. 2019. Cephalopod biodiversity of the Kermadec Islands: implications for conservation and some future taxonomic priorities. Invertebrate Systematics, 33(2): 402–425. doi: 10.1071/IS18041

Carlini, D.B., Young, R.E., Vecchione, M. 2001. A molecular phylogeny of the octopoda (Mollusca: Cephalopoda) evaluated in light of morphological evidence. Molecular Phylogeny and Evolution, 21(3): 388–397.

Chun, C., 1903. Aus den Tiefen des Weltmeeres (2nd edition). G. Fischer, Jena. 592pp.

Clarke, M.R. 1986. A Hand Book for the Identification of Cephalopod Beaks. Clarendon Press, Oxford. 273pp.

Healy, J.M. 1990. Ultrastructure of spermiogenesis in *Vampyroteuthis infernalis* Chun—a relict cephalopod mollusk. Helgoländer Meeresuntersuchungen, 44: 95–107. doi: 10.1007/BF02365433

Hoving, H.T., Robison, B.H. 2012. Vampire squid: detritivores in the oxygen minimum z2one. Proceedings Of The Royal Society B-Biological Sciences, 279: 4559–4567. doi: 10.1098/rspb.2012.1357

Jereb, P., Roper, C.F.E., Norman, M.D., Julian, K.F. (eds) 2010. *Cephalopods of the World. An Annotated and Illustrated Catalogue of Cephalopod Species Known to Date. Volume 3. Octopods and Vampire Squids. FAO Species Catalogue for Fishery Purpose*s, 4(3). Rome, FAO. 353pp.

Joubin, L. 1912. Etudes preliminaires sur les Cephalopodes recueillis au cours des croisieres de S.A.S. le Prince de Monaco. 1re Note: Melanoteuthis lucens nov. gen. et ap. *Bulletin de L’Institut Oceanographique*, 220: 1–14.

Joubin, L.1929. Notes preliminaires sur les Cephalopodes des Croisieres du Dana (1921-22). Octopodes, 1re partie. *Annales de L’Institut Oceanographique*, 6(4): 363–394.

Joubin, L. 1931. Notes preliminaries sur les cephalopodes des Croissieres du Dana (1921-1922). 3e partie. *Annales de L’Institut Oceanographique*, 10(7): 167–211.

Lindgren, A., Giribet, G., Nishiguchi, M. 2004. A combined approach to the phylogeny of Cephalopoda (Mollusca). Cladistics, 20: 454–486. doi: 10.1111/j.1096-0031.2004.00032.x

Lu, C.C., Ickeringill, R. 2002. Cephalopod beak identification and biomass estimation techniques: tools for dietary studies of southern Australian finfishes. Museum Victoria Science Reports, 6: 1–65.

Norman, M.D., Nabhitabhata, J., Lu, C.C. 2016. An updated checklist of the cephalopods of the South China Sea. Raffles Bulletin Of Zoology, 34: 566–592.

Pickford, G.E. 1939. A re-examination of the types of *Melanoteuthis lucens* Joubin. *Bulletin de l’Institut Oceanographique, Monaco*, 777: 1–12.

Pickford, G.E. 1940. The Vampyromorpha, living fossil Cephalopoda. Transactions of the New York Academy of Sciences, 2: 169–181.

Pickford, G.E. 1946. *Vampyroteuthis infernalis* Chun an archaic dibranchiate cephalopod I. Natural history and distribution. Dana-Report, 29: 1–39.

Pickford, G.E. 1949. *Vampyroteuthis infernalis* Chun an archaic dibranchiate cephalopod II. External anatomy. Dana-Report, 32: 1–132.

Robson, G.C. 1932. A Monograph of the Recent Cephalopoda: Based on the Collections in the British Museum (Natural History). Part II, Octopoda. British Museum of Natural History, London. 359pp.

Roper, C.F.E., Voss, G.L. 1983. Guidelines for taxonomic descriptions of cephalopod species. Memoirs of the National Museum of Victoria, 44: 48–63.

Qiu, D., Huang, L., Liu, S., Lin, S. 2011. Nuclear, mitochondrial and plastid gene phylogenies of *Dinophysis miles* (Dinophyceae): Evidence of variable types of chloroplasts. PLoS One, 6(12): e29398. doi: 10.1371/journal.pone.0029398

Sasaki, M. 1920. Report of Cephalopods collected during 1906 by the United States Bureau of Fisheries Steamer “Albatross” in the Northwestern Pacific. Proceedings of the United States National Museum, 57(2310): 163–203.

Schwarz, R., Piatkowski, U., Robison, B.H., Laptikhovsky, V.V., Hoving, H.J. 2020. Life history traits of the deep-sea pelagic cephalopods *Japetella diaphana* and *Vampyroteuthis infernalis*. Deep-Sea Research Part I: Oceanographic Research Papers, 164: 103365. doi: 10.1016/j.dsr.2020.103365

Timm, L.E., Bracken-Grissom, H.D., Sosnowski, A., Breitbart, M., Vecchione, M., Judkins, H. 2020. Population genomics of three deep-sea cephalopod species reveals connectivity between the Gulf of Mexico and northwestern Atlantic Ocean. Deep-Sea Research Part I: Oceanographic Research Papers, 158: 103222. doi:10.1016/j.dsr.2020.103222

Yokobori, S.I., Lindsay, D.J., Yoshida, M., Tsuchiya, K., Yamagishi, A., Maruyama, T., Oshima, T. 2007. Mitochondrial genome structure and evolution in the living fossil vampire squid, *Vampyroteuthis infernalis*, and extant cephalopods. Molecular Phylogenetics and Evolution, 44(2): 898–910. doi: 10.1016/j.ympev.2007.05.009

Young, J.Z. 1977. Brain, behaviour and evolution in cephalopods. *In*: Nixon, M., Messenger, J.M. (eds). The Biology of Cephalopods. Academic Press, New York. pp. 377–434.

Young, R.E. 1972. The systematic and areal distribution of pelagic cephalopods from the seas off southern California. Smithsonian Contributions to Zoology, 97: 1–159.

Young, R.E. 2019. Vampyroteuthidae Thiele, in Chun, 1915. *Vampyroteuthis infernalis* Chun, 1903. The Vampire Squid. Available from http://tolweb.org/Vampyroteuthis_infernalis/20084/2019.03.26 (accessed 22 April 2024)

Young, R.E., Vecchione, M. 1996. Analysis of morphology to determine primary sister-taxon relationships within coleoid cephalopods. American Malacological Bulletin, 12: 91–112.

Young, R.E., Vecchione, M., Donovan, D.T. 1998. The evolution of coleoid cephalopods and their present biodiversity and ecology. In: Payne, A.I.L., Lipinski, M.R., Clarke, M.R., Roeleveld, M.A.C. (eds). Cephalopod Biodiversity, Ecology and Evolution. National Book Printers, Cape Town. pp. 393–420.

